# Genomic features and evolution of the conditionally dispensable chromosome in the tangerine pathotype of *Alternaria alternata*

**DOI:** 10.1101/207746

**Authors:** Mingshuang Wang, Huilan Fu, Xing-Xing Shen, Ruoxin Ruan, Nicholas Pun, Jianping Xu, Hongye Li, Antonis Rokas

**Author notes:** Corresponding authors: Hongye Li and Antonis Rokas, E-mail address (H. Li); (A. Rokas).

## Abstract

The tangerine pathotype of the ascomycete fungus *Alternaria alternata* is the causal agent of citrus brown spot, which can result in significant losses of both yield and marketability for tangerines and tangerine hybrids worldwide. A conditionally dispensable chromosome (CDC), which harbors the host-selective ACT toxin gene cluster, is required for tangerine pathogenicity of *A. alternata*. To understand the genetic makeup and evolution of the tangerine pathotype CDC, we analyzed the function and evolution of the CDC genes present in the *A. alternata* Z7 strain. The 1.84Mb long CDC contains 512 predicted protein-coding genes, which are enriched in functional categories associated with ‘metabolic process’ (132 genes, p-value = 0.00192) including ‘oxidation-reduction process’ (48 genes, p-value = 0.00021) and ‘lipid metabolic process’ (11 genes, p-value = 0.04591). Relatively few of the CDC genes can be classified as CAZymes (13), kinases (3) and transporters (20). Differential transcriptome analysis of H_2_O_2_ treatment and control conditions revealed that 29 CDC genes were significantly up-regulated and 14 were significantly down-regulated, suggesting that CDC genes may play a role in coping with oxidative stress. Evolutionary analysis of the 512 CDC proteins showed that their evolutionary conservation tends to be restricted within the genus *Alternaria* and that the CDC genes evolve faster than genes in the essential chromosomes. Interestingly, phylogenetic analysis suggested that the genes of 13 enzymes and one sugar transporter residing in the CDC were likely horizontally transferred from distantly related species. Among these, one carboxylesterase gene was transferred from bacteria but functionally knocking out this gene revealed no obvious biological role. Another 4 genes might have been transferred from *Colletotrichum* (Sordariomycetes) and 5 were likely transferred as a physically linked cluster of genes from *Cryptococcus* (Basidiomycota) or *Penicillium* (Eurotiomycetes). Functionally knocking out the 5-gene cluster resulted in an 80% decrease in asexual spore production in the deletion mutant. These results provide new insights into the function and evolution of CDC genes in *Alternaria*.

**Author Summary:** Many fungal phytopathogens harbor conditionally dispensable chromosomes (CDCs). CDCs are variable in size, contain many genes involved in virulence, but their evolution remains obscure. In this study, we investigated the origin of the CDC present in the tangerine pathotype of *Alternaria alternata* Z7 strain. We found that most of the Z7 CDC proteins are highly conserved within the genus *Alternaria* but poorly conserved outside the genus. We also discovered that a small number of genes originated via horizontal gene transfer (HGT) from distantly related fungi and bacteria. These horizontally transferred genes include a carboxylesterase gene that was likely acquired from bacteria, a cluster of 4 physically linked genes likely transferred from *Colletotrichum*, and a cluster of 5 physically linked genes likely transferred from *Cryptococcus* (Basidiomycota) or *Penicillium* (Eurotiomycetes). To gain insight into the functions of these transferred genes, we knocked out the bacterial carboxylesterase and the 5-gene cluster. Whereas the carboxylesterase deletion mutant showed no obvious phenotype, the 5-gene cluster mutant showed a dramatically reduced production of asexual spores (conidia). The results of our study suggest that *Alternaria* CDCs are largely comprised from rapidly evolving native genes; although only a few genes were acquired via horizontal gene transfer, some of them appear to be involved in functions critical to the phytopathogenic lifestyle.

## Introduction

*Alternaria alternata* fungi can be ubiquitously found in soil, various plants, and decaying plant debris (Thomma, 2003). Some *A. alternata* strains, which are known as pathotypes, can cause plant diseases and result in severe crop losses worldwide. Specifically, at least seven pathogenic *A. alternata* pathotypes, each producing a unique host-selective toxin (HST) essential to pathogenicity, have been recognized to cause diseases in Japanese pear, strawberry, tangerine, apple, tomato, rough lemon and tobacco (Tsuge et al., 2013). Generally, genes required for HST biosynthesis are clustered on relatively small chromosomes of 1.0 ~ 2.0 Mb in size in *A. alternata* (Tsuge et al., 2013). These chromosomes, which are known as accessory or conditionally dispensable chromosomes (CDCs), are highly variable among species and are required for pathogenicity in *A. alternata*. However, CDCs are generally not required for fungal growth and reproduction on artificial media (Johnson et al., 2001; Hatta et al., 2002).

The importance of CDCs conferring pathogenicity to fruits has been demonstrated by the construction of *A. alternata* hybrids in laboratory conditions. More specifically, two distinct laboratory hybrids were constructed from tomato and strawberry pathotypes and separately for apple and tomato pathotypes. The resulting hybrids harbored two CDCs from their parents allowing for the production of HSTs that caused diseases in both parental plant species (Akamatsu et al., 2001; Akagi et al., 2009b). Furthermore, these studies support the hypothesis that CDCs can be transmitted between different stains thereby facilitating the spread and evolutionary diversification of fungal phytopathogens.

Besides *A. alternata*, many other fungal phytopathogens have CDCs in their genomes including *Fusarium oxysporum, Nectria haematococca, Zymoseptoria tritici* (previously known as *Mycosphaerella graminicola* and *Septoria tritici*), and *Colletotrichum gloeosporioides* (Coleman et al., 2009; Ma et al., 2010; Stukenbrock et al., 2010). Because CDCs are typically found in some, but not all, strains, they have been proposed to have different evolutionary origins than the essential chromosomes (Covert, 1998; Tsuge et al., 2013), e.g., through horizontal transfer from distantly related species (Covert, 1998; Hatta et al., 2002). The possibility of horizontal transfer of CDCs between species has been demonstrated in several studies. For example, a 2-Mb chromosome was transferred between two different biotypes of *C. gloeosporioides* during vegetative co-cultivation in the laboratory (He et al., 1998). Importantly, CDC acquisition can facilitate the transition from the non-pathogenic to pathogenic phenotype in the recipient organism. For example, co-incubation of non-pathogenic and pathogenic genotypes of *F. oxysporum* revealed that the transfer of a CDC from the pathogenic (donor) to non-pathogenic genotype (recipient) enabled the recipient to become pathogenic to tomatoes (Ma et al., 2010).

Previously, the ~1.0 Mb CDC of the tomato pathotype of *A. alternata* was identified and characterized (Hu et al., 2012). Genes in that CDC are abundant in the categories of “metabolic process” and “biosynthetic process”, and include 36 110 polyketide and non-ribosomal peptide synthetase domain-containing genes; furthermore, the GC content of third codon positions, codon usage bias, and repeat region load in the CDC are different from those in the essential chromosomes (Hu et al., 2012), raising the hypothesis that the *A. arborescens* CDC was acquired through horizontal transfer from an unrelated fungus (Hu et al., 2012).

The tangerine pathotype of *A. alternata*, which can cause citrus brown spot on tangerines and tangerine hybrids, was demonstrated to harbour an additional chromosome of about 1.9~2.0Mb by pulse-field gel electrophoresis studies (Miyamoto et al., 2008; Miyamoto et al., 2009). Seven genes, *ACTTR, ACTT2, ACTT3, ACTT5, ACTT6, ACTTS2* and *ACTTS3*, were located onto this chromosome and found to be required for the biosynthesis of the ACT-toxin, a unique HST produced by the tangerine pathotype (Tsuge et al., 2013). However, relatively little is known about the content, potential biological functions, and evolution of the other genes in the CDC of the tangerine pathotype of *A. alternata*. To address this question, we examined the functional annotation, transcriptional activity, and evolution of all 512 genes in the CDC of the tangerine pathotype strain Z7 of *A. alternata*. We found that these genes are enriched in functions associated with ‘metabolic process’. Furthermore, 43 CDC genes were differentially expressed in response to oxidative stress. Finally, we found that conservation for the majority of the 512 Z7 CDC genes was restricted to the genus *Alternaria* and that 14 CDC genes likely originated via horizontal gene transfer (HGT) events from other fungi or bacteria. Construction of the deletion mutant of a carboxylesterase gene that was likely acquired from bacteria did not show any obvious differences in phenotype from the wild type. In contrast, the deletion mutant of a 5-gene cluster that was likely acquired from distantly related fungi showed a dramatic reduction in the production of asexual spores. These results suggest that CDC genes in *Alternaria* are involved in processes associated with metabolism, that they are rapidly evolving, and that they sometimes originate via HGT.

## Results

### General features of CDC

To identify the genome content of CDC of the tangerine pathotype of *A. alternata* strain Z7, we compared the genome sequence of the Z7 strain to that of *A. brassicicola*, which is known to not have a CDC (see methods). This strategy identified 43 CDC contigs with a combined total length of 1.84Mb, which is close to the CDC size estimated by the pulse-field electrophoresis experiment (1.9~2.0Mb) (Miyamoto et al., 2008; Miyamoto et al., 2009). The overall G+C content of the CDC was 47.7%, while that of the essential chromosomes (ECs) was 51.2%. The percentage of repetitive sequences on the Z7 CDC was 1.23%, over 2-fold of that of ECs (0.51%). The average gene length and gene density of CDC were significantly smaller than those of ECs (Table 1).

**Table 1.**
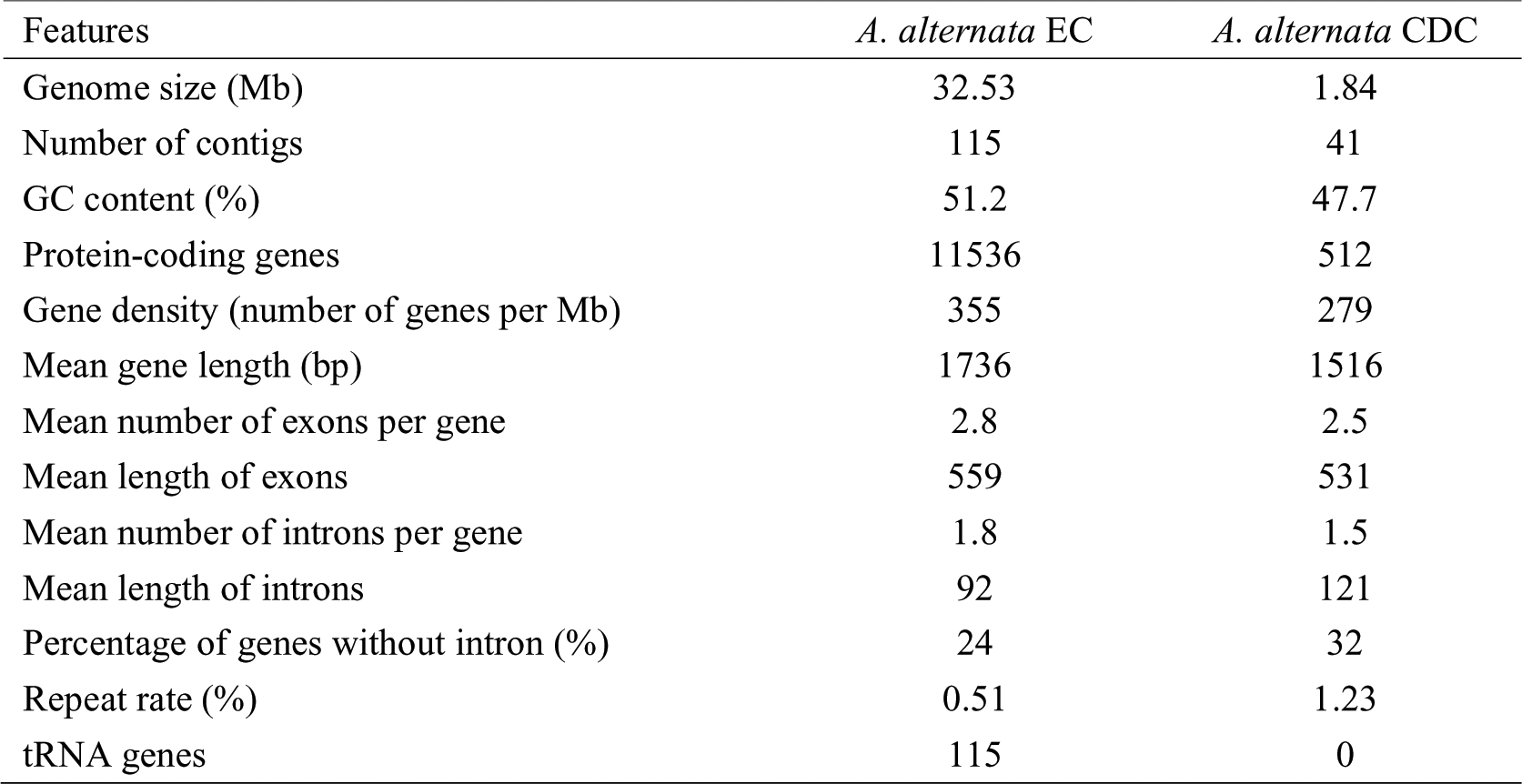
General features of the essential chromosomes (EC) and conditionally dispensable chromosome (CDC) in *A.alternata* strain Z7

### Functional annotation of the genes in the CDC

The CDC was predicted to comprise of 512 protein-coding genes. To predict their functions, gene ontology analysis was performed and 233 genes were assigned to 154 gene ontology terms related to biological process. 132 genes are assigned to the GO term metabolic process (p-value 0.00192) (Fig 1). Within the metabolic process, the GO terms oxidation-reduction process (48, p-value 0.00021) and lipid metabolic process (11, p-value 0.04591) were also significantly enriched (Fig 1). The reduction-oxidation (redox)-associated genes include monoxygenase, dehydrogenase, and reductase, suggesting that the CDC may be involved in intracellular redox homeostasis.

**Fig 1.**
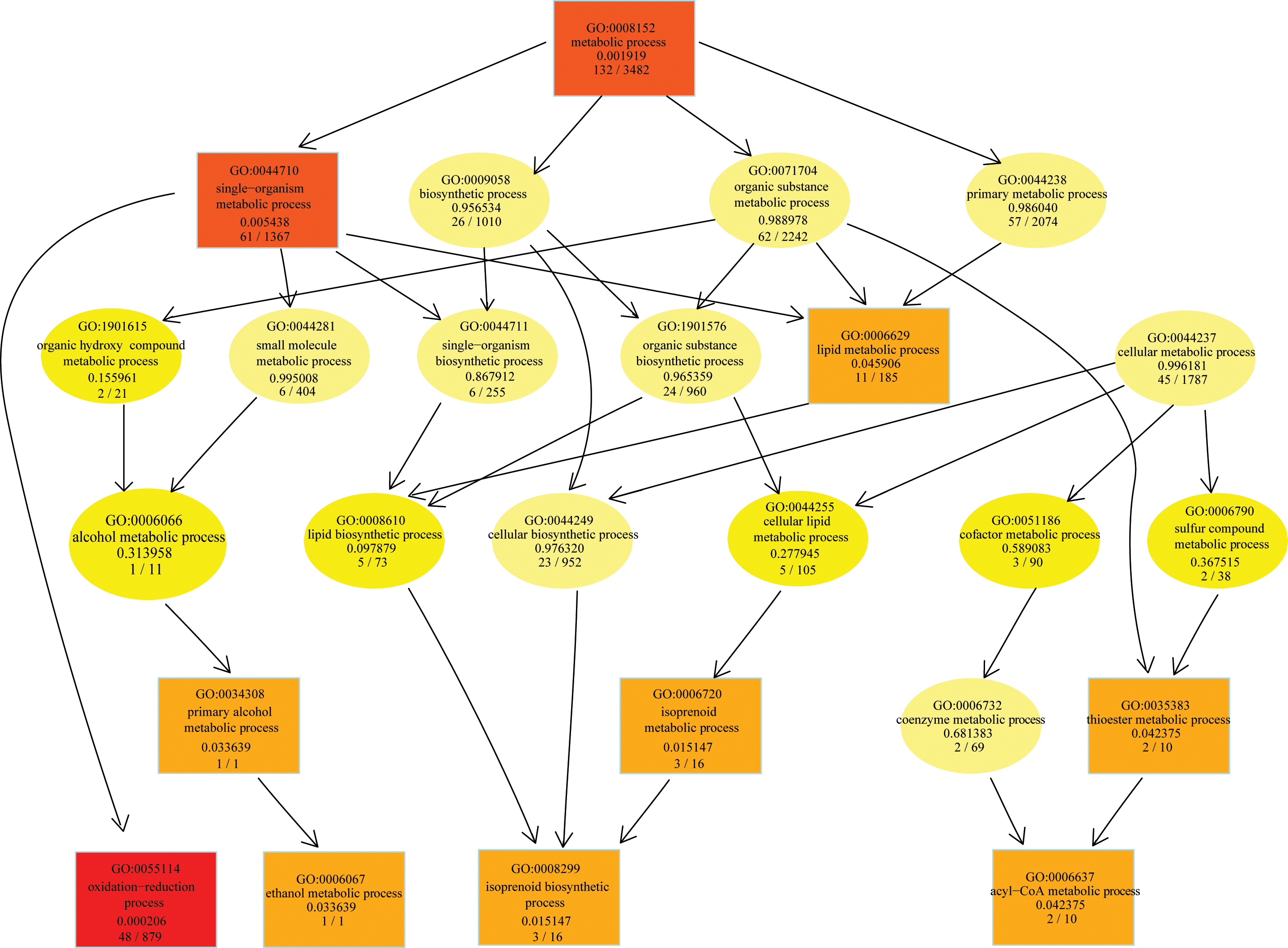
GO enrichment analysis of the Z7 CDC genes in the category “Biological Process”. Significantly enriched GO terms (p<0.05) are illustrated by rectangles. Rectangle color corresponds to degree of statistical significance and ranges from bright yellow (least significant) to dark red (most significant). For each node, the GO identifier, the GO term name, and p-value of each functional category is shown. The final line inside each node shows the number of genes that belong to the functional category in the CDC and in the whole genome of *A. alternata* Z7, respectively.

To functionally annotate the 512 genes in the CDC, we classified their protein products into protein families using several different approaches. Based on Pfam domain characterization, 307 / 512 genes belonged to 195 protein families (S1 Table). We identified 13 CAZyme genes in the CDC, accounting for 3.48% (13 / 373) of the total CAZymes of Z7. These include 3 Glycosyl Transferases (GTs), 6 Glycoside Hydrolases (GHs) and 5 Auxiliary Activities (AAs) (S1 Table). A total of 29 secreted proteins were predicted in the CDC, including 3 plant cell wall-degrading enzymes, 10 small-secreted cysteine-rich proteins (SSCPs), 3 peptidases, 1 lipase, and 1 LysM domain-containing protein (S1 Table). Only 3 kinases from 3 different kinase families were found in the CDC (S1 Table). We identified 20 transcription factors in the CDC, which can be divided into 5 subfamilies: Zinc finger Zn_2_-Cys_6_ (8), Zinc finger C_2_H_2_ (5), Myb-like DNA-binding (1), helix-turn-helix, Psq (4) and high mobility group box (2) (S1 Table). Twenty-six transporter-encoding genes were found in the CDC (S1 Table). Among the 48 redox related genes in the CDC, 13 were predicted to be Cytochrome P450 monooxygenases. Based on comparisons with proteins in the PHI-180 database, 35 CDC proteins are related to pathogenicity (S1 Table). Finally, the CDC contained the ACT-toxin biosynthetic gene cluster present only in the tangerine pathotype of *A. alternata*, which has been described in detail previously (Wang et al., 2016).

### Expression of CDC genes under H_2_O_2_ stress

The production of the host-selective ACT toxin is crucial for the pathogenicity of the tangerine pathotype of *A. alternata* (Miyamoto et al., 2008; Miyamoto et al., 2009; Tsuge et al., 2013). Additionally, recent studies have shown that the ability to eliminate ROS by the tangerine pathotype of *A. alternata* is also of vital importance for pathogenesis to citrus (Lin et al., 2009; Chen et al., 2013; Yang et al., 2016). To discover which genes in the *A. alternata* Z7 CDC are potentially involved in coping with oxidative stress, we performed transcriptome analysis using the previously published transcriptome data of *A. alternata* Z7 after H_2_O_2_ treatment using no H_2_O_2_ treatment as a control (Wang et al., 2016).

Of the 512 CDC genes, 43 were significantly differentially expressed during the H_2_O_2_ stress condition (Table 2, S2 Table). These included 29 CDC genes that were significantly upregulated and 14 that were significantly downregulated. The set of upregulated genes includes proteins such as oxidoreductases, hydrolases, transcription factors and phosphatases. Interestingly, the polyketide synthase (AALT_g11750, log2FC 3.3), which was hypothesized to be crucial for the biosynthesis of the ACT toxin (Miyamoto et al., 2010), was strongly induced after H_2_O_2_ treatment. A cluster of four genes (AALT_g11772, AALT_g11773, AALT_g11774 and AALT_g11775) located on CDC was also highly upregulated (log2FC from 3.4 to 4.4) during H_2_O_2_ stress. After comparing these four proteins to the Pfam database, proteins AALT_g11772 and AALT_g11774 were found to contain an oxidoreductase family domain and a NmrA-like family domain, respectively. However, no protein domain was predicted for AALT_g11773 and AALT_g11775 (Table 2, S2 Table).

**Table 2.**
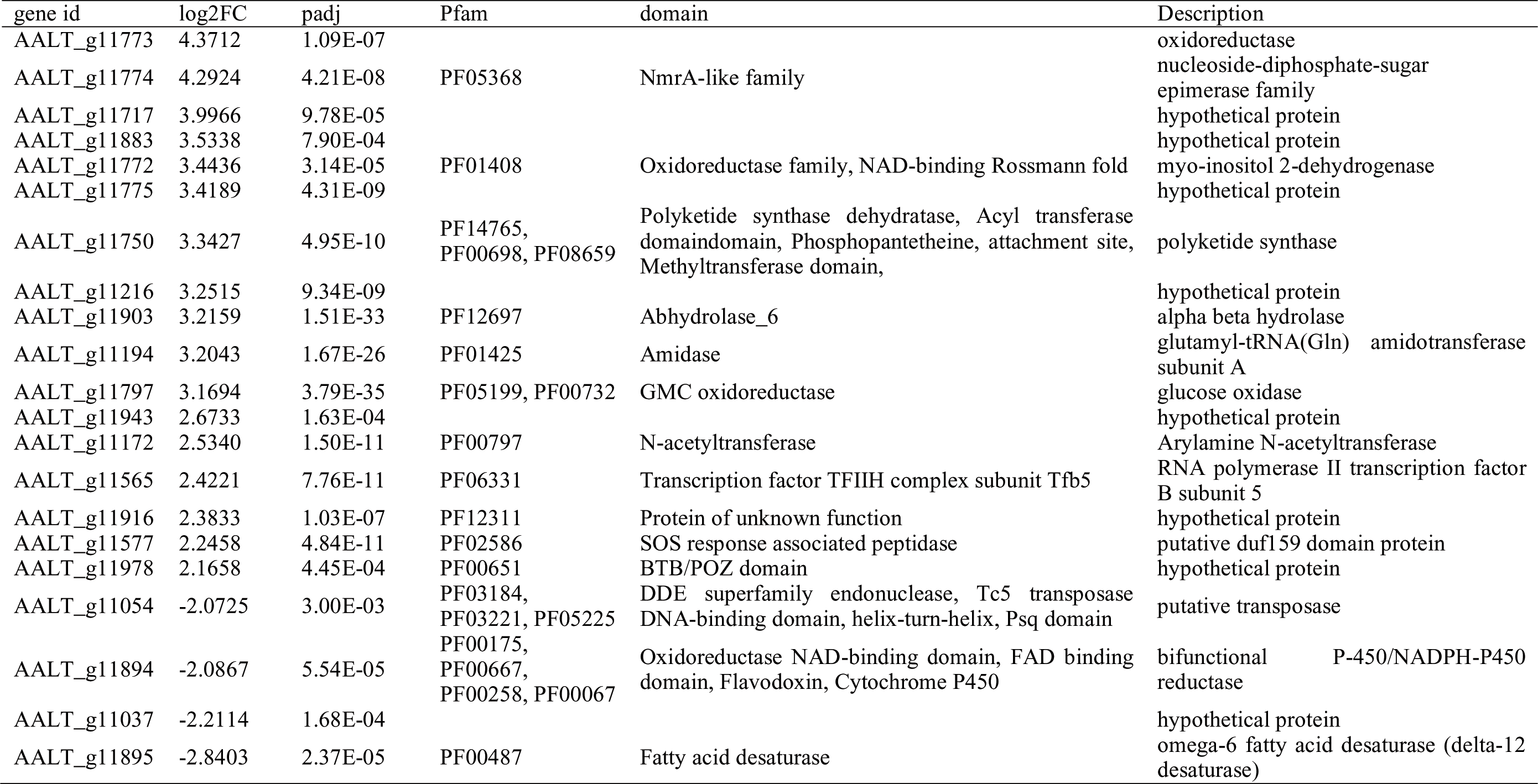
Annotation of the differentially expressed genes located in the conditionally dispensable chromosome with absolute log2FC >2 during H_2_O_2_ stress

### The Evolutionary origin of the Z7 CDC

To explore the evolutionary origin of the Z7 CDC, we compared the sequence 2 similarity (calculated by BLAST identity score * query coverage) of each of the 512 2 CDC proteins to the protein sequences from the genomes of other species in the 2 Dothideomycetes. In the species phylogeny, *Alternaria* species are grouped into three 2 clades (I through III), which coincides with a previously constructed phylogeny based 2 on 200 conserved single-copy orthologs (Wang et al., 2016). Proteins in the Z7 CDC 2 showed a wide range of sequence similarity values to proteins in other 2 Dothideomycetes (Fig 2). As expected, the highest degree of similarity was with 2 proteins of other *Alternaria* species, and in particular with those from clade I (Fig 2, 2 S1 Fig). For example, we found that 442 / 512 (86.3%) of the Z7 CDC proteins (442, 86.3%) showed >50% similarity to proteins in *A. turkisafria* (Fig 2, S1 Fig), which can also cause citrus brown spot. However, *A. citriarbusti* and *A. tangelonis* can also cause citrus brown spot but these exhibited much lower numbers (321 / 512, 62.7% and 310 / 512, 60.5%, respectively) of proteins with >50% similarity to proteins in Z7 CDC (Fig 2). These results suggest that the gene content and sequence similarity on CDC can be highly variable among strains of the same pathotype.

**Fig 2.**
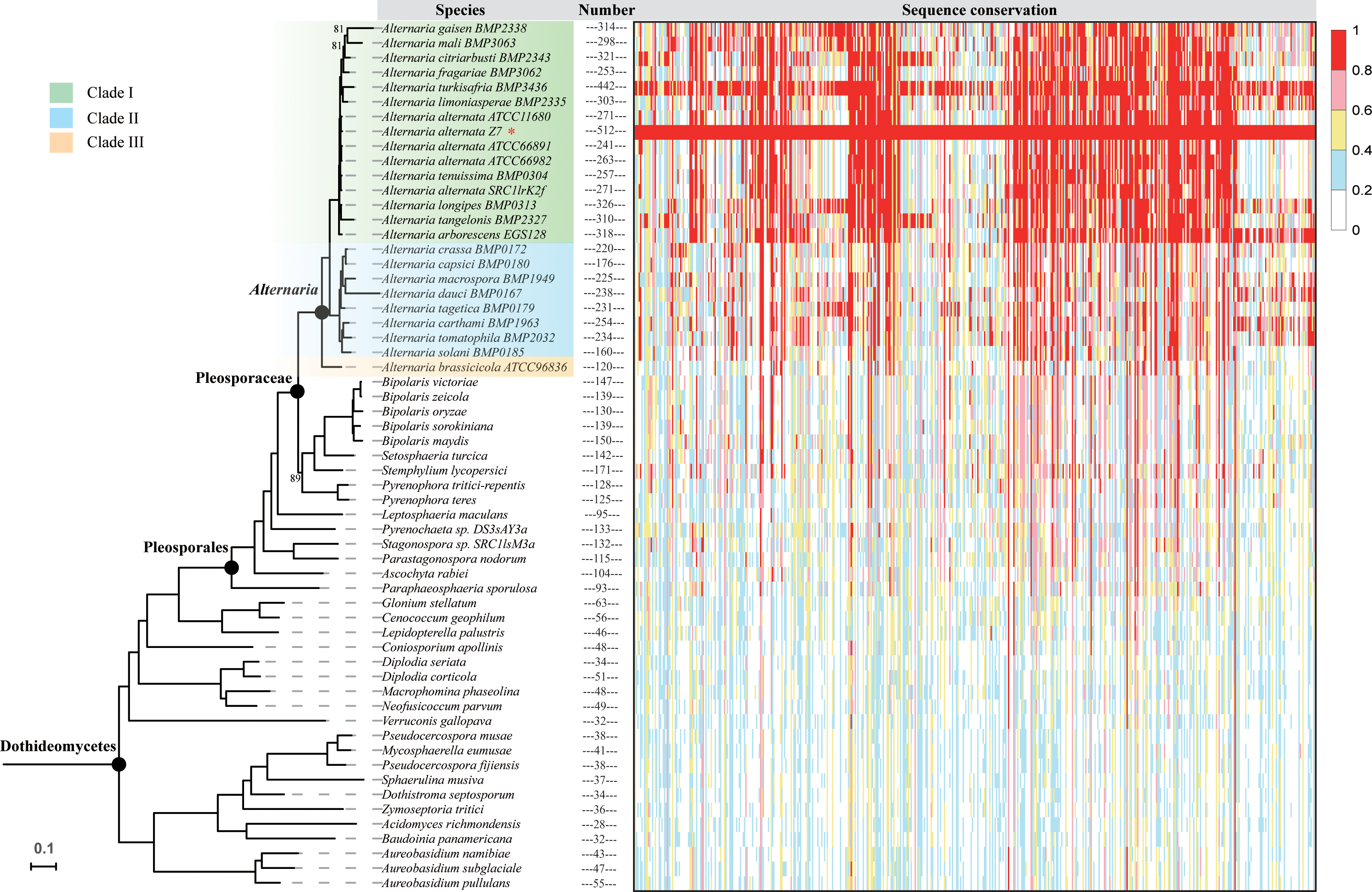
Sequence conservation of Z7 CDC genes across the phylogeny of *Alternaria* and representative species of Dothideomycetes. The phylogeny on the left depicts the evolutionary relationships of species of *Alternaria* and representative Dothideomycetes. The maximum likelihood phylogeny was inferred from the concatenation-based analysis of an amino acid data matrix comprised from 1,754 single-copy BUSCO genes under the LG+R10 substitution model. Branches with bootstrap support values of 100% are not shown; branches with bootstrap values <100% are shown near each branch. The heat map on the right was constructed using the sequence conservation value (BLAST identity * query coverage) of each Z7 CDC protein to its best counterpart in each of the species included in this analysis. Cells with red color correspond to proteins (in specific species) that exhibit high sequence conservation to a given Z7 CDC protein. The numbers next to each species’ name correspond to the number of proteins that exhibit sequence conservation values greater or equal to 0.5 when compared to Z7 CDC proteins.

To examine if any of the Z7 CDC proteins were more similar to proteins outside those found in genomes from the genus *Alternaria*, a BLASTp search of those 512 230 protein sequences against the NCBI non-redundant database was performed and the result was filtered using the E-value<1e-10 and sequence identity>30% criteria. Although the best matches for 330 / 512 proteins were proteins from other *Alternaria* species, 81 proteins had their best matches to be proteins found in non-*Alternaria* members from the family Pleosporaceae, 33 from the order Pleosporales (other than Pleosporaceae), 9 from the class Dothideomycetes (other than Pleosporales), 44 from the domain Fungi (other than Dothideomycetes), and 1 from the domain Bacteria (S3 Table).

To further dissect the evolutionary history of Z7 CDC genes, we reconstructed the phylogenetic tree for each Z7 CDC gene with their orthologs from other species in the Dothideomycetes. There were too few orthologs to construct phylogenetic trees for three CDC genes; among the remaining 509 gene trees, 402 showed monophyly within the genus *Alternaria*. Taken together, our results suggest that most of the *A.alternata* Z7 CDC proteins were likely present in the *Alternaria* ancestor and that some of them were likely independently lost in some of the species during the diversification of the genus.

### Relative evolutionary rate of Z7 CDC

To figure out if genes in the CDC and genes in the ECs in Z7 evolve differently, we built a phylogenetic tree for each Z7 gene which has orthologs in more than 7 251 (50% of total number) species within Alternaria Clade I. A total of 191 CDC and 10,060 EC genes were used to calculate the average branch length for each tree. The average branch length for most genes in both the CDC and the ECs are very low (Fig 3), which indicates that most genes are highly conserved within the genus *Alternaria* Clade I. However, as a whole, we found that EC genes had lower average branch lengths than CDC genes (P= 2.2e-16, Fig 3), suggesting that the Z7 CDC genes are evolving faster than the Z7 EC genes.

**Fig 3.**
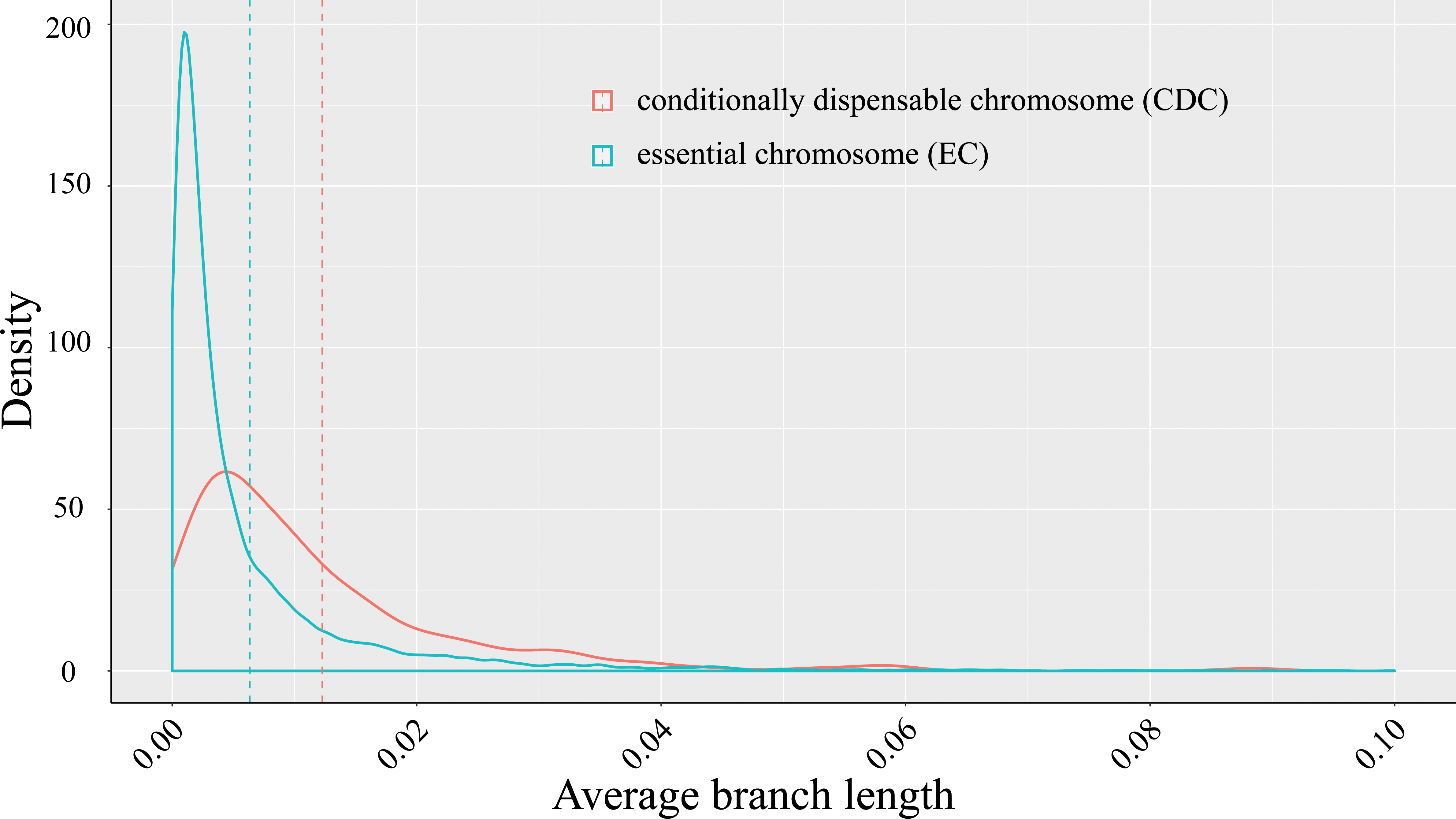
Distribution of the average branch length from all individual Z7 CDC (red line) or EC (blue line) gene trees constructed from groups of orthologous genes within *Alternaria* Clade I. The dashed lines denote the mean values of the two distributions.

### HGT of genes in the CDC

To examine whether any of the *A. alternata* Z7 CDC genes originated via HGT, we calculated the Alien Index (Gladyshev et al., 2008; Wisecaver et al., 2016) of all 512 genes. A total of 43 genes show AI > 0 and at least 80% of their top 200 BLASTp hits with a taxonomic classification other than Dothideomycetes. The validity of these 43 HGT candidates was further examined phylogenetically. The phylogenetic trees for most of these 43 HGT candidates were weakly supported, but the evolutionary origin of 14 of these genes was strongly supported to be outside Dothideomycetes (Table 3, S2-15 Fig). The approximately unbiased (AU) test for each of the 14 genes significantly rejected the hypothesis that they formed a monophyletic group with the rest of the sequences from Dothideomycetes (Table 3). As the genes inferred to have undergone HGT are also found in other *Alternaria* species, we infer that the HGT events occurred before the divergence of the Z7 strain from the other *Alternaria* genomes examined and not after. Specifically, 4 of the horizontally transferred genes are found in the genomes of species in *Alternaria* Clades I and II, 6 genes are also found in the genomes of species in *Alternaria* Clade I, and 4 genes are found only in the Japanese pear, strawberry and tangerine pathotypes (Table 3).

**Table 3.**
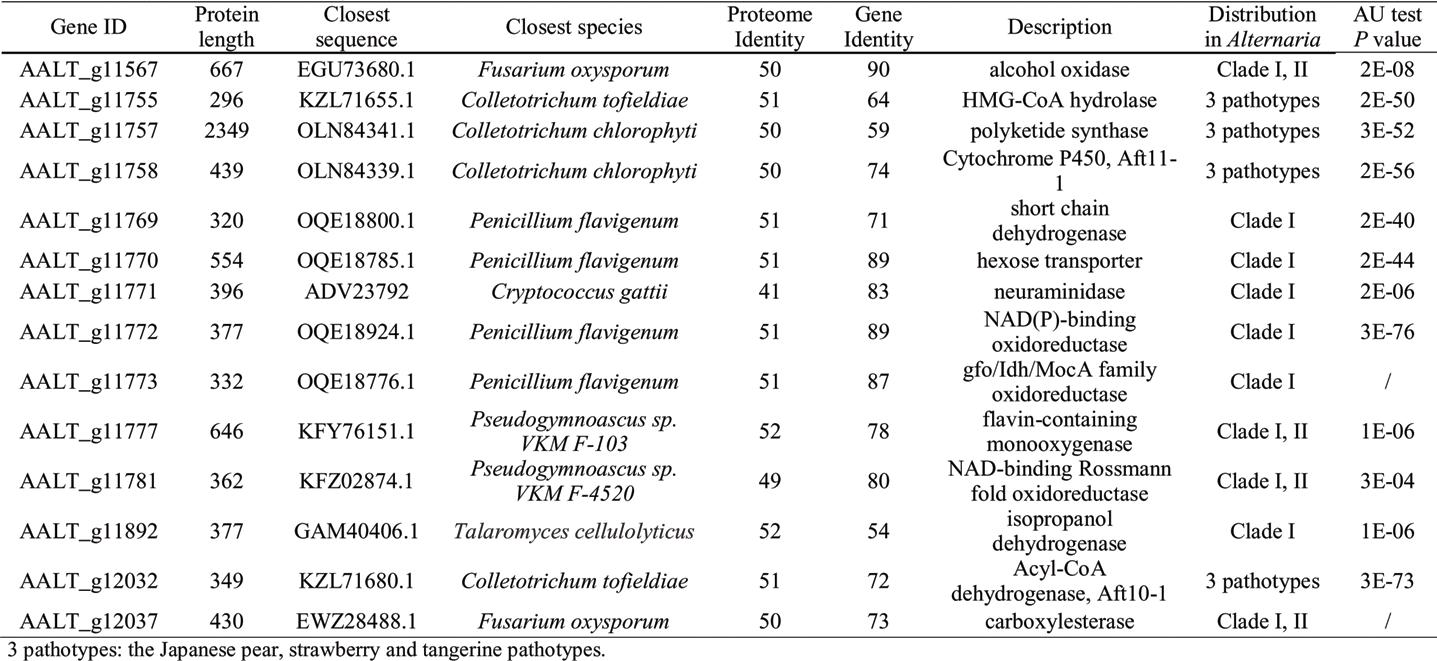
Summary list of all horizontal gene transfers (HGTs) in *A.alternata* Z7 CDC.

All HGT genes encode enzymes except for AALT_g11770, which is a hexose transporter (Table 3). According to their predicted functions, these HGT candidates are involved in two biological processes: oxidation reduction and carbohydrate metabolism (Table 3). There is only one gene (AALT_g12037) that was likely transferred from bacteria (Fig 4A, Fig S2). The phylogenetic tree of this gene contains a large proportion of bacteria (Fig 4A). There are only 15 fungal species predicted to contain this gene, including 13 *Alternaria* species, *Fusarium oxysporum* and *Pyrenochaeta sp. DS3sAY3a* (Fig 4B). AALT_g12037 is 1,428 bp in length and contains no introns. It encodes one of the carboxylesterases that are responsible for the hydrolysis of carboxylic acid esters into their corresponding acid and alcohol (Potter and Wadkins, 2006). This protein in Z7 CDC shows high sequence identity (75%, 73% and 74%) and query coverage (99%, 100% and 98%) to its orthologs in *Fusarium oxysporum, Pyrenochaeta sp. DS3sAY3a* and *Bacillus subtilis*, one of the potential donors. To test the functional significance of this gene, the coding region of AALT_g12037 was knocked out in *A.alternata Z7* (Fig S16A, B). However, the mutant strain did not exhibit any phenotypic difference from the wildtype strain (Fig S16E, F, G).

**Fig 4.**
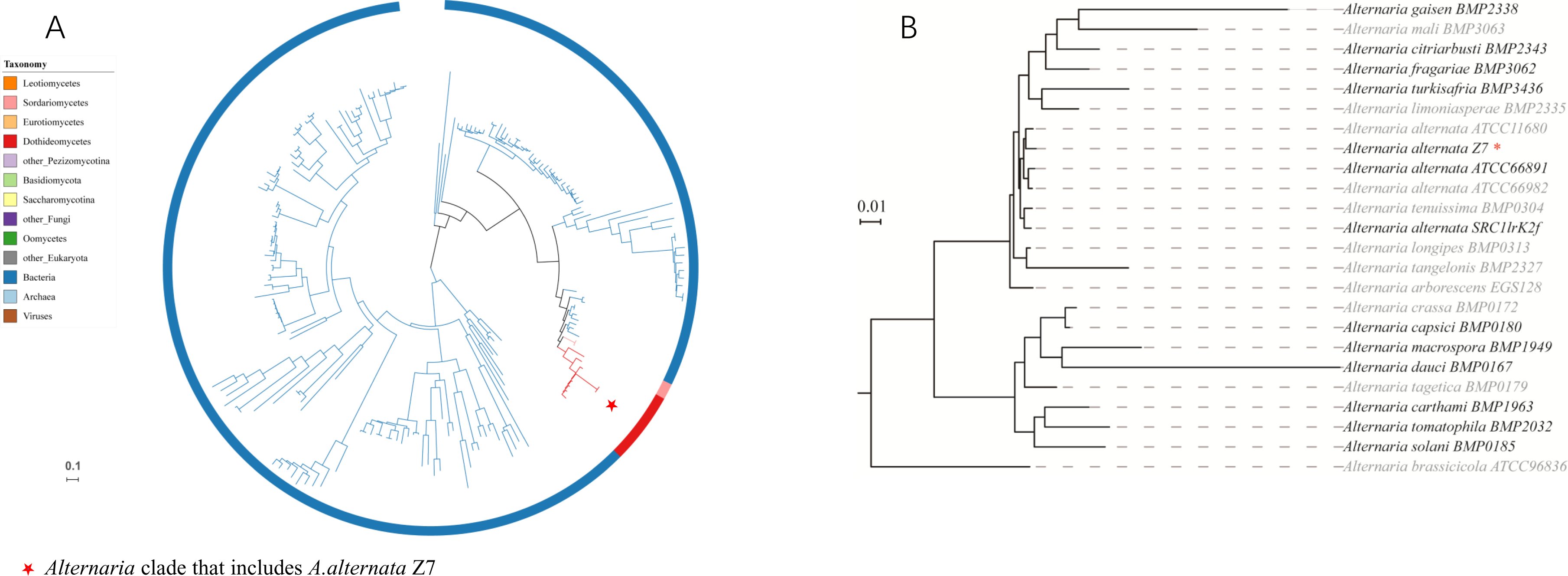
The gene AALT_g12037 was horizontally transferred from Bacteria to the *A. alternata* Z7 CDC. A) ML phylogeny of AALT_g12037 homologs across the tree of life. The gene’s maximum likelihood phylogeny was inferred under the best substitution model automatically selected by ModelFinder, as implemented in IQ-TREE 1.5.4. Branch colors indicate the taxonomic lineages to which the different taxa included in each phylogeny belong. The red asterisk indicates the *Alternaria* clade on each phylogeny. The full tree can be found in S2 Fig. B) The presence of the HGT-acquired gene family in other *Alternaria* genomes. Light grey indicates the lack of this HGT gene in the corresponding species.

The ACT toxin is essential for the pathogenicity of *A. alternata* Z7 to citrus leaves, the synthesis of which is predicted to be controlled by a cluster composed of about 25 genes (Wang et al., 2016). Surprisingly, 4 / 25 of the ACT cluster genes contained in the Z7 CDC strain are always grouped together with sequences from the unrelated *Colletotrichum* (Sordariomycetes) in their gene phylogenies (Fig 5A, S3-6 Fig). Interestingly, the orthologous genes in *Colletotrichum tofieldiae* are physically linked with each other and are part of a secondary metabolite biosynthetic gene cluster predicted by antiSMASH 4.0 (Blin et al., 2017) (Fig 5A, B), although the gene order and orientation of the two clusters is different (Fig 5B). Besides *A.alternata* Z7 (the tangerine pathotype), this cluster is also present in the Japanese pear pathotype and the strawberry pathotype (Wang et al., 2016). Previously, deletion of the AKT3 gene in the Japanese pear pathotype, which is the ortholog of the HMG-CoA hydrolase gene ACTT3 (AALT_g11755), produced toxin-deficient and non-pathogenic mutants (Tanaka and Tsuge, 2000). Taken together, these results raise the hypothesis that the HGT of four genes from a lineage related to *Colletotrichum* may have contributed to the composition of the HST gene clusters found in the CDCs of three pathotypes of *Alternaria*.

**Fig 5.**
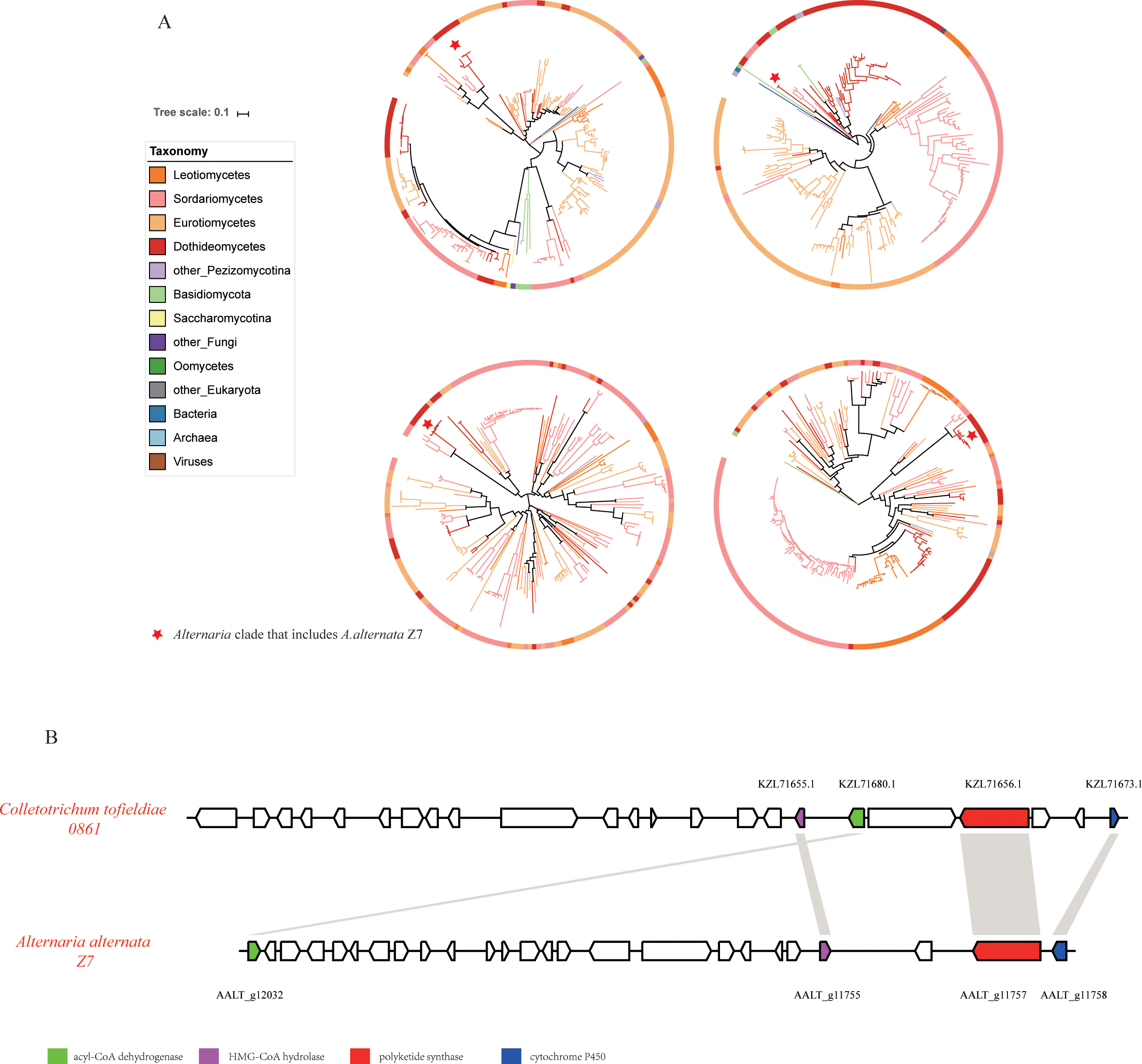
Horizontal transfer of 4 genes in the ACT toxin gene cluster in the tangerine pathotype of *A.alternata* Z7 CDC. A) Phylogenetic evidence of the HGT of these genes. For each gene, the maximum likelihood phylogeny was inferred under the best substitution model automatically selected by ModelFinder, as implemented in IQ TREE 1.5.4. Branch colors indicate the taxonomic lineages to which the different taxa included in each phylogeny belong. The red asterisk indicates the *Alternaria* clade on each phylogeny. The full phylogenetic trees of the individual genes can be found in S3 – S6 Figs. B) Conservation of synteny among the ACT toxin gene cluster in *A.alternata* Z7 and a predicted secondary metabolic gene cluster in *Colletotrichum tofieldiae* 0861. Orthologs among different species are marked with same color and homologous regions are shown by the gray boxes. Arrows indicate gene direction.

Interestingly, five of the transferred genes are physically clustered in the Z7 strain CDC and almost always appear on the gene phylogeny as sisters to sequences from either *Cryptococcus* (Basidiomycota) or *Penicillium* (Eurotiomycetes) fungi (Fig 6A, S7-11 Fig). Specifically, both *Penicillium flavigenum* and *Cryptococcus gattii* have 4 317 clustered genes that are highly similar and group together on the gene phylogeny with 4 of these 5 genes (Fig 6A, B); *Penicillium flavigenum* lacks the neuraminidase encoding gene homolog (AALT_g11771) while *Cryptococcus gattii* lacks the oxidoreductase encoding gene homolog (AALT_g11772) (Fig 6B). However, the gene order and orientation is quite different among the clusters of these three genomes (Fig 6B). From these results, the most likely scenario is that the genes were horizontally transferred from *Cryptococcus* or *Penicillium* species and were subsequently rearranged. To further characterize this 5-gene cluster, we constructed a deletion mutant lacking the entire cluster and examined its phenotypic characteristics upon osmotic, oxidative, fungicide, and cell wall stresses, as well as its asexual development and pathogenicity (S16 Fig). We found that the mycelial growth of the mutant in potato dextrose agar (PDA) was slightly reduced (by 9.1%) when compared with that of the wildtype (S16G Fig); furthermore, the mutant formed a fluffy colony during asexual development and conidial production was decreased by 79.5% relative to wild type conidial production) (Fig 6C). No other phenotypic differences between the mutant and the wildtype strains were found (S16E, G Fig). These results indicate that this horizontally transferred gene cluster is critical for the conidiation of *A. alternata*.

**Fig 6.**
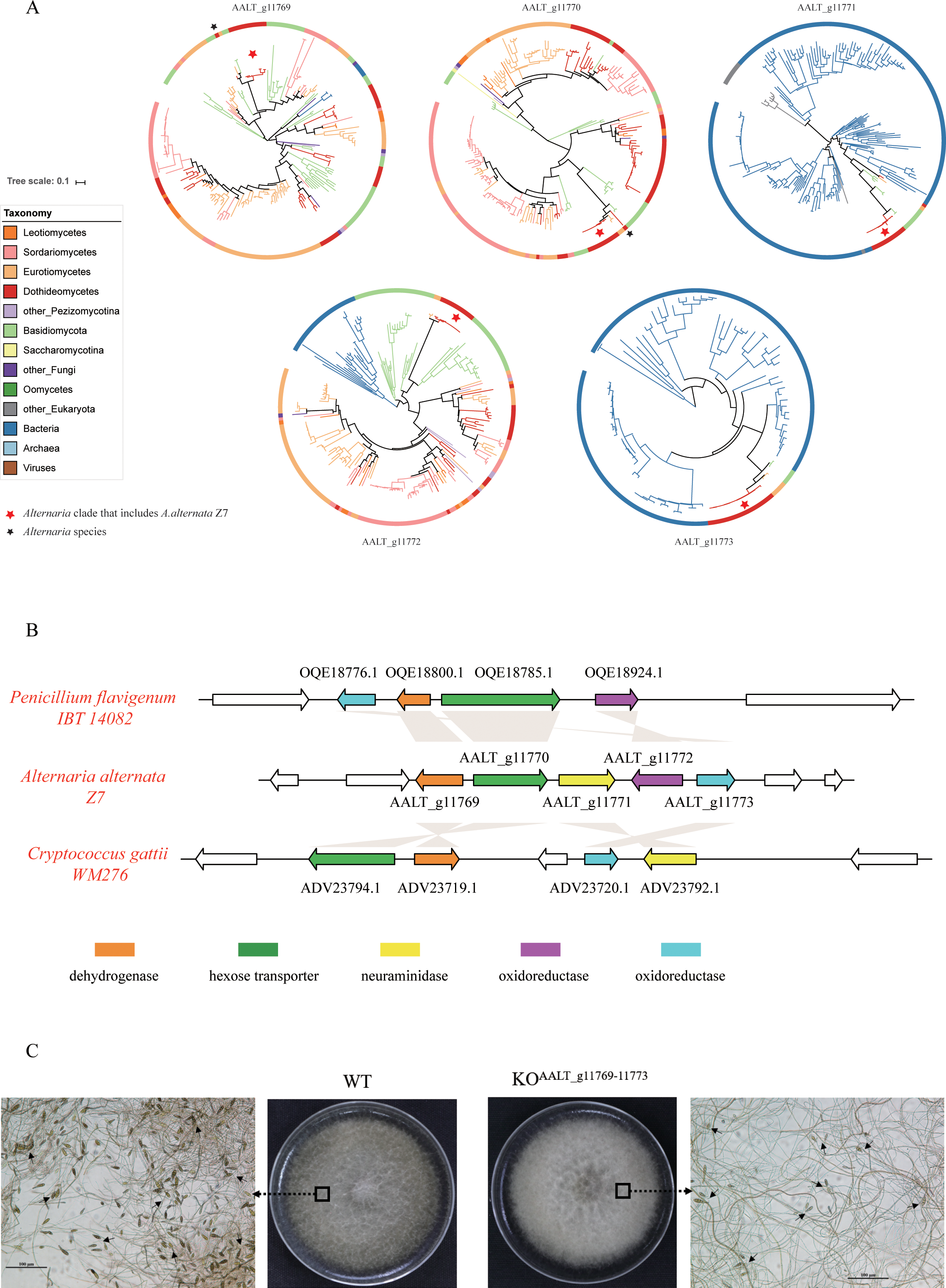
Horizontal transfer of a cluster of 5 genes in the tangerine pathotype of *A.alternata* Z7 CDC. A) Phylogenetic evidence of the HGT of these genes. For each gene, the maximum likelihood phylogeny was inferred under the best substitution model automatically selected by ModelFinder, as implemented in IQ-TREE 1.5.4. Branch colors indicate the taxonomic lineages to which the different taxa included in each phylogeny belong. The red asterisk indicates the *Alternaria* clade on each phylogeny. The full phylogenetic trees of the individual genes can be found in S7 S11 Figs. B) Conservation of synteny among the cluster of 5 genes in *A. alternata* and the evolutionary related clusters present in *Penicillium flavigenum* and *Cryptococcus gatti*. Orthologs among different species are marked with same color and homologous regions are shown by the gray boxes. Arrows indicate gene direction. C) Conidial production in the wildtype (WT) and in the 5-gene cluster deletion mutant (KO^AALT_g11679-11773^) after incubation for 7 days. Arrows indicate representative conidia (not all of the conidia are indicated). Bar, 100µm.

## Discussion

Conditionally dispensable chromosomes (CDCs) are commonly found in fungal phytopathogens and play key roles in pathogenicity (Johnson et al., 2001; Hatta et al., 2002). However, despite their importance, our knowledge about the gene content and evolutionary origin of CDCs is limited. In this study, we identified and characterized the 1.84 Mb CDC of the tangerine pathotype strain Z7 of *A. alternata* and examined the function and evolutionary history of its genes. Our results suggest that CDC genes in *Alternaria* are involved in processes associated with metabolism, that they are conserved within the genus *Alternaria* and are rapidly evolving, and that they sometimes originate via HGT. Below, we discuss our findings in the context of the genome content of *Alternaria* CDCs and the evolutionary origin of *Alternaria* CDC genes.

In this study, we functionally annotated the CDC of the tangerine pathotype strain Z7 of *A. alternata*. Although the Z7 CDC is much larger than the *A. arborescens* CDC, several Z7 CDC gene families are comparable to those present in the *A. arborescens* CDC, including CAZymes, SSCPs, kinases, transcription factors and transporters (S1 Table) (Hu et al., 2012). This result suggests that there are some shared properties between CDCs from these two divergent species. Our analyses also identified differences between CDCs in *A. arborescens* and in *A. alternata* strain Z7. For example, the enrichment of “biosynthetic process” genes was discovered in the *A. arborescens* CDC but not in the *A. alternata* CDC (Hu et al., 2012). In addition, the *A. arborescens* CDC contains 10 SM clusters and harbors 36 polyketide and non-ribosomal peptide synthetase genes that might be responsible for the biosynthesis of the backbone structures of several groups of secondary metabolites (Hu et al., 2012). However, in Z7 CDC, except for the host selective ACT-toxin gene cluster, no other gene cluster involved in the biosynthesis of secondary metabolites was identified. About 39.2% (201) of the 512 genes found in Z7 CDC were described as hypothetical proteins according to their BLAST hits in the NCBI nr database, indicating that the functions of a great number of Z7 CDC genes are not known (S1 Table).

Evolutionary analysis showed that most of the Z7 CDC proteins are highly conserved within the genus *Alternaria* and likely originated in the *Alternaria* last common ancestor (Fig 2). These results contrast with those of a previous study suggesting that the *A. arborescens* CDC originated from an unknown species through HGT (Hu et al., 2012). The discrepancy is most likely due to the small amount of available data used in the previous study; specifically, *A. arborescens* protein-coding genes were compared to only two small databases respectively composed of either only *A. brassicicola* proteins or of three fungal species from other filamentous fungal genera (*Leptosphaeria maculans, Pyrenophora tritici-repentis*, and *Aspergillus oryzae*) (Hu et al., 2012). In contrast, the much larger number of *Alternaria* genomes examined in our study shows that the evolutionary history of most CDC genes is consistent with the species phylogeny and with vertical, not horizontal, transmission.

Comparison of the Z7 CDC proteins to those in the proteomes of other *Alternaria* species showed that CDCs in different species or even in the same pathotype can be very variable (Fig 2). This result is consistent with the fact that the length of CDCs in distinct pathotypes of *A.alternata* is very variable. For example, both the strawberry and the tomato pathotypes contain a ~1.0 Mb CDC (Ito et al., 2004; Hu et al., 2012); both the Japanese pear and the tangerine pathotypes contain a 1.9-2.0 Mb CDC (Miyamoto et al., 2008; Miyamoto et al., 2009); the apple pathotype contains a 1.1-1.8 Mb CDC (Harimoto et al., 2007); while the rough lemon pathotype contains a 1.2-1.5 Mb CDC (Masunaka et al., 2005). Although the mechanism underlying the formation of these diverse CDCs in *Alternaria* species is largely unknown, a recent population genomics study in *Zymoseptoria tritici* identified the precise breakpoint locations of insertions that give rise to the highly differentiated gene contents in *Z. tritici* CDCs (Croll et al., 2013). That same study also reported the occurrence of CDC losses in progeny because of nondisjunction during meiosis as well as the emergence of a new CDC through a fusion between sister chromatids (Croll et al., 2013). Thus, the CDCs of *Z. tritici* were proposed to originate mainly from ancient core chromosomes through a degeneration process involving breakage-fusion-bridge cycles and large insertions (Croll et al., 2013). Although determining whether the breakage-fusion-bridge cycles model holds for *Alternaria* CDCs will require further sequencing of CDCs and analyses, we note that our finding of evolutionary conservation of CDC genes within *Alternaria* is consistent with the *Z. tritici* model.

Horizontal transfer of CDCs between different strains of the same or closely related fungal species has been documented for *Fusarium oxysporum, Colletotrichum gloeosporioides*, as well as for some *Alternaria* species (He et al., 1998; Akamatsu et al., 2001; Akagi et al., 2009a; Ma et al., 2010). Horizontal transfer of entire CDCs may contribute not only to the introduction of CDCs in fungal populations but also to their subsequent spread and to their acquisition of virulence factors that are essential for pathogenicity. Since fungal CDCs are known to be able to transfer between closely related species, it may be very difficult to distinguish between vertical and horizontal transmission of CDCs within *Alternaria*. On the other hand, horizontal transfer of CDCs between distantly related species has never been demonstrated, making it a less likely explanation for the evolutionary origin of *Alternaria* CDCs.

Although horizontal transfer is an unlikely explanation for the formation of CDCs in *Alternaria*, our study found evidence that HGT is a mechanism for the origin of some of the genes residing in its CDCs. One of the Z7 CDC genes, a carboxylesterase, was likely acquired from Bacteria (Fig 4A). Carboxylesterases are ubiquitous enzymes that exist in almost all living organisms and whose function is to hydrolyze carboxylesters into the corresponding carboxylic acid and alcohol (Satoh and Hosokawa, 1998). By degrading exogenous xenobiotics that contain esters, carboxylesterases are thought to be associated with detoxification (Hatfield et al., 2016), although, to date, no endogenous carboxylesterase substrates have been identified. However, functional analysis of the horizontally transferred carboxylesterase did not reveal any difference between a mutant strain lacking the gene and the wild type strain for a range of phenotypes (Fig S16).

The 14 genes that were likely horizontally transferred in the Z7 CDC from distantly related species, included 9 genes that formed 2 gene clusters (Table 3). Previously, the 23-gene secondary metabolic gene cluster involved in the biosynthesis of the mycotoxin sterigmatocystin was shown to have been horizontally transferred from *Aspergillus* to *Podospora* (Slot and Rokas, 2011). HGT of intact gene clusters would not only contribute to fungal metabolic diversity but also potentially provide its recipient with a competitive advantage offered by the ability to synthesize a novel secondary metabolite; for example, a subsequent study showed that *Podospora* produces sterigmatocystin (Matasyoh et al., 2011). Although the horizontally transferred gene cluster in this study contains fewer genes, the 5-gene cluster was shown to be critical for conidial production (Fig 6C). Additionally, the HMG-CoA hydrolase coding gene AKT3 found in another transferred cluster was shown to be absolutely required for the HST production and virulence to host plant (Tanaka and Tsuge, 2000). These results are consistent with the view that HGT events, including ones involving the transfer of entire clusters, have played important roles over the course of the evolution of filamentous fungi (Fitzpatrick, 2012; Soanes and Richards, 2014; Wisecaver and Rokas, 2015).

## Methods

### Genomic and transcriptomic data retrieval

The assembled *A. alternata* Z7 genome and proteome were downloaded from GenBank under the accession number LPVP00000000 (Wang et al., 2016). Genome data from other *Alternaria* species were downloaded from the *Alternaria* genomes database (Dang et al., 2015) and from other species of Dothideomycetes from the GenBank database (S4 Table) (last access on April 15, 2017). The transcriptome data of *A. alternata* after H_2_O_2_ treatment were downloaded from the NCBI’s Sequence Read Archive (SRA) database with accession number SRP071688 (last access on May 8, 2017).

### Identification of the contigs comprising the CDC

To identify all the contigs that are part of the *A. alternata* Z7 CDC, we used a previously described method with slight modifications (Hu et al., 2012). Briefly, all *A. alternata* Z7 contigs with sequence lengths greater than 5 kb were aligned to contigs from *Alternaria brassicicola*, a species that belongs to the same genus as *A. alternata* but which does not carry CDCs, using MUMmer 3.0 with an identity cut-off at 80% (Delcher et al., 2003). All contigs whose sequence coverage when aligned to *A. brassicicola* was less than 20% were considered to be parts of the *A. alternata* Z7 CDC.

### Functional annotation of the genes in the CDC

To functionally annotate the 512 genes (and their protein products) in the CDC, we performed gene ontology analysis using the topGO version 2.28.0 (Alexa and Rahnenfuhrer, 2016) and classified the 512 proteins into protein families using the Pfam, version 31.0 databases (last access on June 1, 2016) (Finn et al., 2014). We predicted CDC genes that were parts of fungal secondary metabolite pathways using the web-based analytical tool SMURF (http://www.jcvi.org/smurf/index.php, last access on Jun 20, 2016) (Khaldi et al., 2010). To identify CDC proteins that are carbohydrate-active enzymes, we searched with the CAZymes Analysis Toolkit (http://www.cazy.org/, last access on Jul 10, 2016) based on sequence and Pfam annotation (Lombard et al., 2014). To identify secreted proteins, we used SignalP, version 4.1 to predict transmembrane domains (Petersen et al., 2011) and we excluded non-extracellular and GPI-anchored proteins by using ProtComp, version 5 (Klee and Ellis, 2005) and fragAnchor (http://navet.ics.hawaii.edu/~fraganchor/NNHMM/NNHMM.html, last access on July 10, 2016) (Poisson et al., 2007), respectively. All resulting secreted proteins that were shorter than 200 amino acids (aa) in length and contained at least 4 cysteine residues were considered as small secreted cysteine-rich proteins. Kinases were searched and classified by an automated pipeline (Kosti et al., 2010). Cytochrome P450s were classified based on Pfam and the P450 database, version 1.2 (last access on June 18, 2016) (Moktali et al., 2012). Transporters were identified by performing BLASTp against the Transporter Classification Database (last access on June 7, 2016) (Saier et al., 2016).

### Species phylogeny inference and evaluation of sequence conservation of CDC genes across Dothideomycetes

To understand the origin and evolution of genes in the *A. alternata* CDC, we first constructed a species phylogeny using genomic data from all available *Alternaria* species as well as for representative species from other Dothideomycetes. We constructed the species phylogeny using 1,754 conserved, fungal BUSCO genes as described previously (Simao et al., 2015; Shen et al., 2016). Briefly, the gene structure of each genome was predicted by AUGUSTUS 3.1 (Stanke and Waack, 2003), then the sequences of these predicted genes were aligned to the HMM alignment profile of each BUSCO gene in the OrthoDB v9 database (Simao et al., 2015), and the ones with alignment bit-scores higher than 90% of the lowest bit-score among the reference genomes were kept for tree construction. The isolates *Alternaria porri BMP0178* and *Alternaria destruens BMP0317* were excluded from downstream analyses due to their poor (<60%) coverage of BUSCO genes (S5 Table). We also compared the sequence similarity of each of the 512 CDC proteins to proteins in the genomes of other Dothideomycetes by calculating the value of BLAST identity * query coverage.

### Examination of relative evolutionary rate between genes in the CDC and genes in the essential chromosomes

To estimate the relative evolutionary rate between Z7 CDC and essential chromosomes (ECs), ortholog groups within the *Alternaria* Clade I containing Z7 proteins from both CDC and ECs were extracted using OrthorMCL v2.0.9 and reciprocal BLASTp with identity>50% and query coverage >50% as cut-offs (Chen et al., 2006). For each ortholog group containing more than 7 (50% of total number) species, only one sequence per isolate/species (the one that was the best hit to the Z7 protein) was kept for further analysis. The coding sequences of those ortholog proteins were then aligned with MAFFT v7.023b using the E-INS-I strategy (Yamada et al., 2016) and trimmed with trimAl v1.4.rev11 using its automated1 strategy (Capella-Gutierrez et al., 2009). The maximum-likelihood (ML) phylogenetic trees were inferred using IQ-TREE 1.5.4, with the best model selected by ModelFinder (Nguyen et al., 2015; Kalyaanamoorthy et al., 2017) and with 1,000 bootstrap replicates. The significance of the difference in the average total branch length of the ML phylogenetic trees of CDC genes against that of the EC genes was determined by Wilcoxon test.

### Identification of CDC genes that underwent HGT

To detect gene candidates that experienced HGT in *A.alternata* Z7 CDC, we first performed a BLASTp search of the local NCBI’s nonredundant protein database (nr, last access on May 21, 2017) using Z7 CDC proteins as queries. We next selected proteins with the following characteristics as HGT candidates for further phylogenetic analyses: a) an Alien Index (AI) score larger than 0 (Gladyshev et al., 2008; Wisecaver et al., 2016), b) at least 80% of the top 200 BLASTp hits of the query protein are from organisms other than Dothideomycetes, and c) the sequence identity of the query protein across its entire length to its best BLASTp hit is equal or greater than 50%.

All genes that fit these three criteria were used as query sequences in BLASTp searches against the nr database and phylogenetic trees of their most closely related sequences across the tree of life were constructed. To reduce the number of sequences used to build each phylogenetic tree, we kept only one sequence per species (the one with the best BLASTp hit to the HGT candidate), then we selected the top 200 hits from the Blast results. The resulting sequences were used as input for multiple sequence alignment, trimming and phylogenetic inference, which were performed as described above.

The phylogenetic tree of each HGT candidate was manually inspected and only those trees that were evidently incongruent with the species phylogeny and strongly supported (bootstrap value > 95%) were retained as HGT candidates. For those HGT candidates, we used the Consel software, version V0.1i (Shimodaira and Hasegawa, 2001; Shimodaira, 2002) to perform the approximately unbiased (AU) comparative topology test between the unconstrained ML tree and the constrained ML tree in which the *Alternaria* gene sequence was forced to be monophyletic with the rest of the sequences from Dothideomycetes. All phylogenetic trees were visualized using ITOL version 3.0 (Letunic and Bork, 2016).

### Functional analyses of horizontally transferred genes

The AALTg12037 gene and the 5-gene cluster were knocked out using a fungal protoplast transformation protocol, as described previously (Chen et al., 2017). Briefly, the two flanking 600-900bp fragments and a bacterial phosphotransferase B gene (HYG) were fused together and the resulting fragment was then introduced into fungal protoplasts using polyethylene glycol and CaCl_2_. The transformants growing on a medium supplemented with 150 μg/ml hygromycin were selected and examined by PCR with specific primer pairs. All the primers used in this study are listed in S6 Table.

Wildtype and mutant strains were grown on regular solid potato dextrose agar (PDA) at 25 ׉ and asexual spore (conidial) production was inspected under a Nikon Eclipse 80i light microscope (Nikon, Tokyo, Japan) after incubating for 7 days. Conidia were collected with 10 mL sterile solution of 0.05% (v/v) Tween 20 and filtered through lens wiping paper. Conidial concentration was quantified using a hemocytometer. To examine stress tolerance, mutant and wildtype strains were grown on PDA plates supplemented with either 1.5mM CuSO4, 0.01% SDS, 250ug/mL Congo red, 250mM CaCl_2_, 1 M NaCl, 25ug/mL Azoxystrobin, 20mM H_2_O_2_ or 2mM tert-Butyl hydroperoxide. Each plate was inoculated with a 5-mm mycelial plug taken from the edge of a 5-day-old colony. The diameters of the colonies were measured after the plates were incubated at 25 °C for 5 days. Fungal virulence was assessed on *Citrus clementina* leaves inoculated by placing a 5mm plug taken from the media for 2 days. Each strain was tested on at least 5 leaves and experiments were repeated two times.

### Data availability

All data generated in this study, including CDC contigs, CDC gene annotation, multiple sequence alignments and phylogenetic trees, have been deposited on the figshare repository at DOI: 10.6084/m9.figshare.5549077 (the data will be made publicly available upon acceptance of the manuscript).

## Acknowledgments

We thank members of the Rokas lab for helpful discussions. We thank Abigail Leavitt Labella and Jacob Steenwyk for their critical comments on this paper. This work was conducted in part using the resources of the Advanced Computing Center for Research and Education (ACCRE) at Vanderbilt University.

## Author contributions

Conceptualization: Mingshuang Wang, Xing-Xing Shen, Antonis Rokas, Hongye Li.

Data curation: Mingshuang Wang.

Formal analysis: Mingshuang Wang.

Funding acquisition: Hongye Li, Mingshuang Wang, Antonis Rokas.

Investigation: Mingshuang Wang, Huilan Fu, Ruoxin Ruan, Nicholas Pun.

Methodology: Mingshuang Wang, Huilan Fu, Xing-Xing Shen, Ruoxin Ruan.

Project administration: Antonis Rokas, Hongye Li.

Resources: Mingshuang Wang.

Software: Mingshuang Wang, Xing-Xing Shen.

Supervision: Antonis Rokas, Hongye Li.

Validation: Mingshuang Wang, Huilan Fu, Ruoxin Ruan.

Visualization: Mingshuang Wang,vHuilan Fu, Xing-Xing Shen,vRuoxin Ruan.

Writing original draft: Mingshuang Wang, Jianping Xu, Hongye Li.

Writing review&editing: Antonis Rokas, Mingshuang Wang.

## Supporting information

S1 Fig. Bar plot of the degree of sequence conservation of Z7 CDC proteins to the proteomes of other species in the Dothideomycetes. The sequence conservation value of each Z7 CDC protein to its best counterpart in each of the species was calculated by BLAST identity * query coverage.

S2 – S15 Figs. Full ML phylogenetic trees of HGT gene homologs across the tree of life. Each gene’s maximum likelihood phylogeny was inferred under the best substitution model automatically selected by ModelFinder, as implemented in IQTREE 1.5.4. Branch colors indicate the taxonomic lineages to which the different taxa included in each phylogeny belong. The *A. alternata* Z7 CDC sequence that was used as a query in the BLAST search is shown in red font. Bootstrap values are shown near each branch. S2 Fig: AALT_g12037; S3 Fig: AALT_g12032; S4 Fig: AALT_g11755; S5 Fig: AALT_g11757; S6 Fig: AALT_g11758; S7 Fig: AALT_g11769; S8 Fig: AALT_g11770; S9 Fig: AALT_g11771; S10 Fig: AALT_g11772; S11 Fig: AALT_g11773; S12 Fig: AALT_g11567; S13 Fig: AALT_g11777; S14 Fig: AALT_g11781; S15 Fig: AALT_g11892.

S16 Fig. Functional analysis of the role of the carboxylesterase gene (AALT_g12037) and horizontally transferred 5-gene cluster (AALT_g11769 – AALT_g11773) of the CDC. A) Schematic depiction of gene disruption of AALT_g12037 via homologous recombination. Numbers denote primers listed in supplementary S6 Table. B) PCR verification of the deletion of the AALT_g12037 gene. Long bands can be only amplified in the mutants (KO^AALT_g12037^) using the outside PCR primer pairs 7/8 while the short band can be only amplified in the wildtype (WT) using the inner PCR primer pair 9/10. C) Schematic depiction of gene disruption within AALT_g11769 – AALT_g11773 via homologous recombination. Numbers denote primers listed in supplementary S6 Table. D) PCR verification of the deletion of AALT_g11769 – AALT_g11773 gene cluster. Long bands can be only amplified in the mutants (KO^AALT_g11679-11773^) using the outside PCR primer pairs 17/18 while the short band can be only amplified in the wildtype (WT) using the inner PCR primer pair 19/20. E) Virulence of the wildtype and mutant strains to citrus leaves. F) Conidial production in the wildtype and mutant strains after incubation for 7 days. Bar, 100µm. G) Growth assays of the wildtype and mutant strains on different conditions. CR: Congo red t-BHP: tert-Butyl hydroperoxide MM: Minimal medium.

